# Evolution and evolvability of rifampicin resistance across the bacterial tree of life

**DOI:** 10.1101/2024.11.05.622190

**Authors:** Negin Bolourchi, Christopher R. P. Brown, Andrew D. Letten, Jan Engelstädter

## Abstract

Predicting the ability of bacteria to develop antibiotic resistance is challenging, especially for the vast majority of species for which no experimental data are available. Here, we investi-gated the evolvability and intrinsic presence of rifampicin resistance across the bacterial tree of life. We compiled a panel of known rifampicin resistance mutations, comprising 60 amino acid substitutions within the gene *rpoB*. We then screened *>* 18 000 genomes from all major bac-terial groups for the presence of those mutations and determined which mutations can evolve through point mutations. Our results demonstrate that although the evolvability of individual mutations varies considerably across species, overall predicted evolvability is high and relatively homogeneous across bacterial taxa. Rifampicin resistance mutations are present intrinsically in 8% of species. Our analysis provides a global picture of the mutational landscape of rifampicin resistance, including both insight into existing observations as well as predictions informing future work.

## Main

The evolvability of a biological system refers to its ability to respond to natural selection^1,2^. Understanding evolvability is essential for predicting adaptive responses to environmental chal-lenges^3,4^. It is hence relevant to a wide range of fundamental and applied problems in biology, but perhaps nowhere more so than in tackling the evolution of antibiotic resistance ^4,5^.

Curtailing antibiotic resistance is predicated on our ability to predict the evolutionary trajectory a microbial lineage is most likely to take^6^. A critical first step towards this goal is to determine the spectrum of resistance mutations available to a given lineage and the number of independent pathways it has to access them. With the exception of the most well studied clinical pathogens and lab strains, whose resistance profiles have been extensively screened (e.g., Refs. 7–11), this information is not readily available for the vast majority of bacteria. Nevertheless, the genomes of known resistant lineages and mutants might still be leveraged to predict the resistance evolvability of understudied taxa.

The mapping of known resistance mutations to unscreened bacteria may also serve to identify taxa that are already intrinsically resistant to certain antibiotics. This is to say we may be able to use current knowledge to make predictions on both the potential to evolve resistance *and* the preexistence of resistance mutations. Given that most antibiotics are derived from compounds produced naturally by bacteria and other microbes^12^, there are likely to be large numbers of species with unrecorded intrinsic resistance to various antibiotics. Identifying taxa that are intrinsically resistant holds promise not just for our fundamental understanding of bacterial diversity but also potentially in the design of probiotics to accompany antibiotic treatment^13^.

In the quest to translate resistance from known to unknown taxa, it is reasonable to expect that the robustness of predictions across taxa will be greatest for antibiotics that 1) target highly conserved sites; 2) cannot be evaded by too large a variety of resistance mechanisms; and 3) whose resistance evolution has already been studied in some depth. One drug that ranks highly for these three criteria is rifampicin (RIF), a broad-spectrum antibiotic that is used to treat a range of bacterial infections and is a first-line drug for the treatment of tuberculosis (caused by *Mycobacterium tuberculosis*)^14,15^. The drug targets the beta subunit of the RNA polymerase, binding close to its active site and blocking elongation of nascent RNA transcripts, thereby leading to premature termination of transcription and ultimately cell death. Resistance to RIF arises predominantly through point mutations in *rpoB*, the gene coding for the beta subunit of the RNA polymerase^15,16^. Prior work in several species has established that almost all RIF resistance mutations arise in one of four, relatively small clusters within *rpoB* : I, II, III and N. Of these, cluster I is the largest and contains most reported resistance mutations, including in clinical isolates of RIF resistant *M. tuberculosis* ^17^.

We set out to predict the evolvability of RIF resistance across the bacterial tree of life. Specifi-cally, using a comprehensive inventory of reported resistance mutations from the literature, we screen the genomes of *>* 18 000 bacterial species spanning all major phyla for the evolvability or pre-existence of RIF resistance. We find that although total evolvability (number of resistance mutations accessible though a single nucleotide change) is largely invariant across species, the spectrum of mutations available varies considerably. In addition, we find that the predicted dis-tribution of existing RIF resistance is clustered phylogenetically, with several clades uniformly possessing known RIF resistance mutations. This includes several groups for which there is direct, but until now unexplained, evidence of RIF resistance.

## Results

### Mutations reported to confer RIF resistance

We compiled data from 116 studies reporting a total of 1426 RIF resistance mutations within the *rpoB* gene (Supplemental Tables S1 and S2). Merging homologous residues reported in different studies and/or species produced a list of 369 distinct single amino acid substitutions at 146 positions. These mutations were reported across 75 bacterial species and included both mutations obtained in experimental studies (mostly mutant screens) and in clinical or envi-ronmental isolates (see Figure 1 for an overview). Mutations 526Y and 526R (all positions in *Escherichia coli* coordinates) had the highest frequencies (reported in 30 and 26 species, respec-tively), and position 526 is also the position with the greatest number of reported mutations (Figure 1A). The largest number of mutations were reported in *Mycobacterium tuberculosis*, *Salmonella enterica*, *E. coli* and *Streptococcus pneumoniae* (Figure 1B).

**Figure 1:**
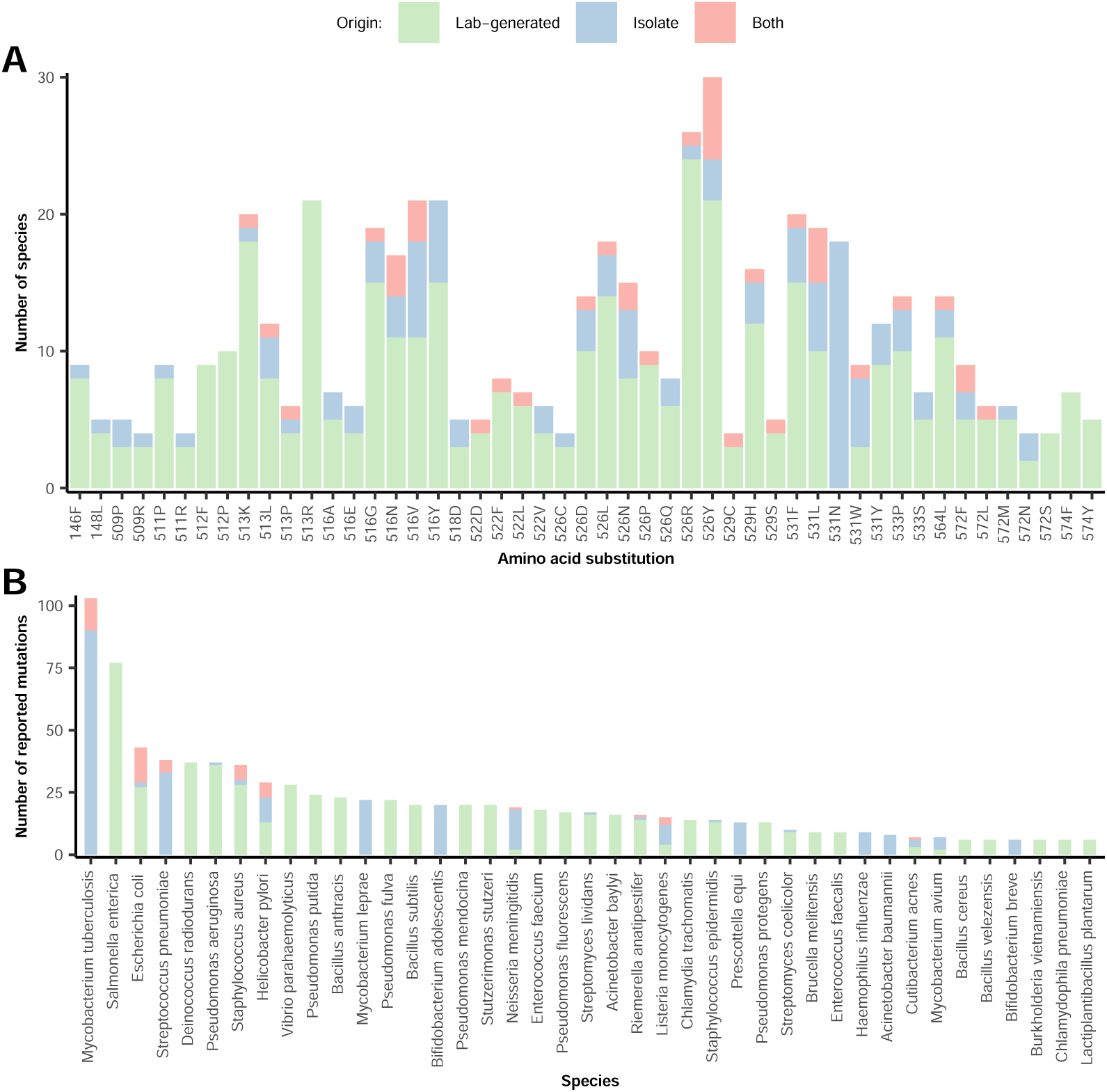
RIF resistance mutations within *rpoB* reported within the literature. A) The number of species in which different resistance mutations were reported, coloured by whether the mutations were generated in experimental studies (mostly mutant screens), in clinical/environmental isolates, or in both settings. Only mutations reported in at least four species are shown. B) The number of mutations reported across species, with the same colour coding as in A. Only species with at least six reported mutations are shown.

From this large pool of reported mutations, we sought to construct a panel of mutations that were likely to robustly confer resistance across species. We therefore discarded mutations that were reported in fewer than three independent studies or three species. After applying these

### The phylogenetic distribution of predicted intrinsic and evolvable RIF resistance across bacteria

We next extracted *rpoB* sequences from 18 469 annotated reference or representative bacterial genomes obtained from the NCBI genome database ^18^. After applying quality-control filters to these sequences, we retained 18 238 high-quality *rpoB* sequences from 18 127 species (Figure S1). (91 species carried more than one *rpoB* copy; see below.) We then screened each sequence for both the presence and evolvability of the 60 mutations in our panel of reported RIF resistance mutations.

Based on this screen, we predict 1453 species (8% of the species in our dataset) to be intrinsi-cally resistant to RIF because they carry at least one of the reported resistance mutations in our panel (Supplemental Table S3). These species are spread across the entire bacterial phylogeny but cluster (permutation test: *p <* 0.001) in distinct bacterial clades (Spirochaetia, Bifidobac-terales, Mollicutes, Erysipelotrichales and others) (Figure 2). Conversely, some large clades (e.g., classes Cytophagia, Sphingobacteriia and Chitinophagia within the phylum Bacteroidea as well as the Enterobacterales) appear to be devoid or almost devoid of any intrinsically resis-tant species.

**Figure 2:**
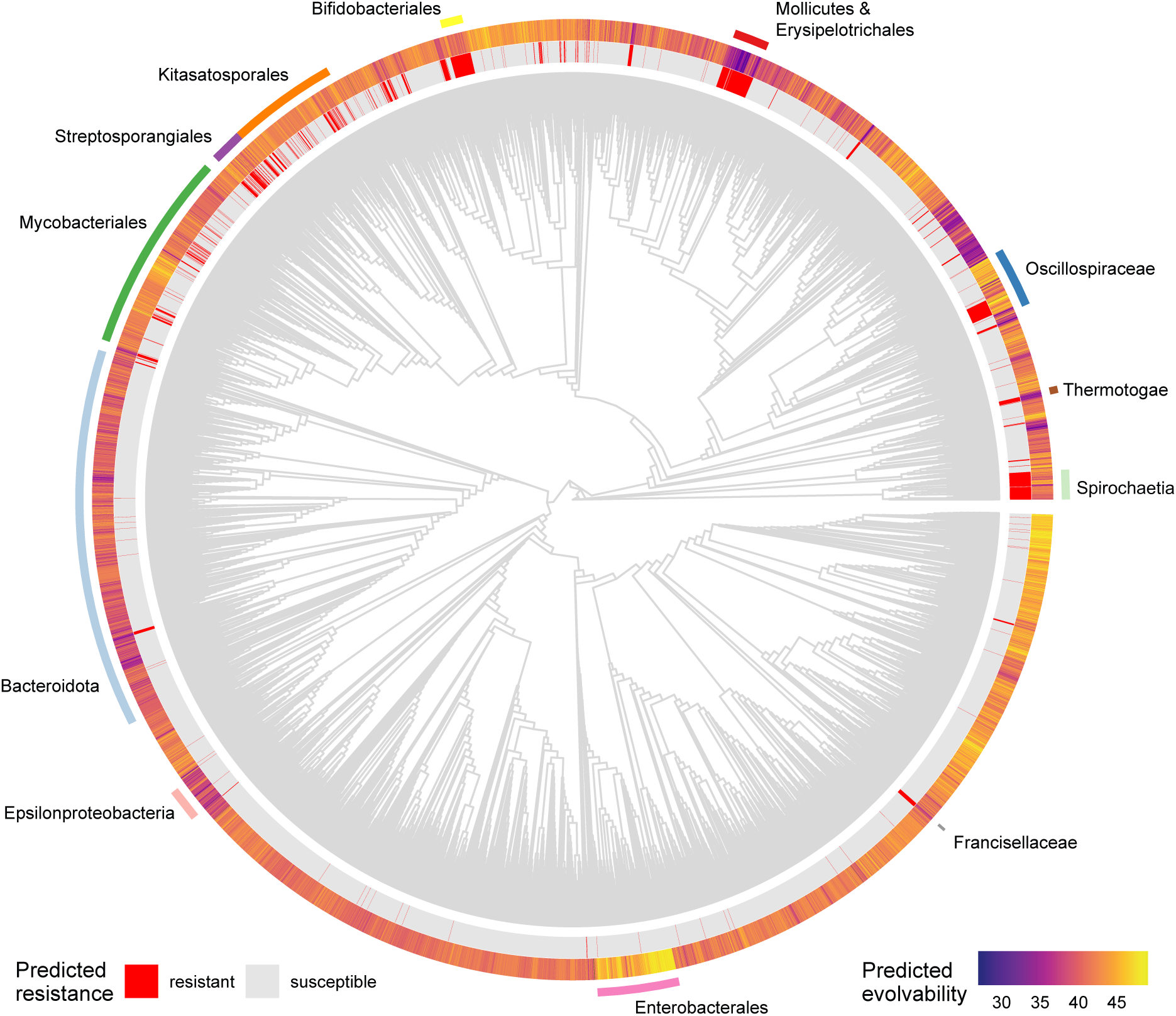
The phylogenetic distribution of predicted intrinsic resistance and evolvability of RIF resistance. The phylogenetic tree is a subtree of the phylogenetic tree available on the Genome Taxonomy Database ^19^, containing the majority (15 521) of our screened species. The inner circle shows predicted intrinsic resistance as red lines (species with at least one amino acid residue from our panel of reported resistance mutations). The outer circle shows the predicted evolvability of RIF resistance (number of amino acid substitutions from our panel that are accessible via a single point mutation).

Evolvability, here defined as the total number of reported RIF resistance mutations (at the amino acid level) that can arise through a single point mutation (at the nucleotide level), varies between 27 and 49 across the bacterial tree of life (95% interquantile range: 35 to 47). The upper limit of this observed range in evolvability is close to the theoretical maximum of 53 (this theoretical maximum, which is considerably below the total number of 60 mutations, arises because many reported mutations are at the same site and are impossible to all evolve from any one codon through a single nucleotide substitution). As observed for intrinsic resistance, there is a clear phylogenetic signal in the distribution of evolvability, with some clades displaying distinctly high or low values (Pagel’s *λ* = 0.74; *p <* 0.001 and Blomberg’s *K* = 0.102; *p* = 0.001).

Figure 3A shows the distribution of evolvability and intrinsic resistance for the largest bac-terial classes. Almost all species within the Spirochaetia carry the 531N mutation and are hence predicted to be intrinsically resistant to RIF. Similarly, almost all members of the Mol-licutes are predicted to be resistant, but in this case due to the presence of 526N (with the exception of 25 species harbouring 526Q). This class also exhibits very low evolvability, indicat-ing that Mollicute species generally cannot easily acquire many of the other known resistance mutations. Other classes comprising many species predicted to be intrinsically RIF resistant include the Actinomycetes (571 species), and the Clostridia (119 species). Aside from well-known genera within the Spirochaetia (e.g., *Borrelia* and *Treponema*) and Mollicutes (e.g., *Spiroplasma* and *Mycoplasma*), unrelated genera such as *Streptomyces* are also predicted to comprise many or even mostly resistant species (Figure 3B). Notably, intrinsic resistance within *Streptomyces* appears to have arisen independently many times as not all members of this genus have RIF resistance mutations despite having a close phylogenetic relationship (Figure S2; see also Discussion). Another, small class that is predicted to be almost universally resistant is the Erysipelotrichia (52 out of 54 species, see also Discussion). A total of 104 species carry two or more resistance mutations, a pattern particularly common in the genera *Nocardia* and *Nonomuraea*.

**Figure 3:**
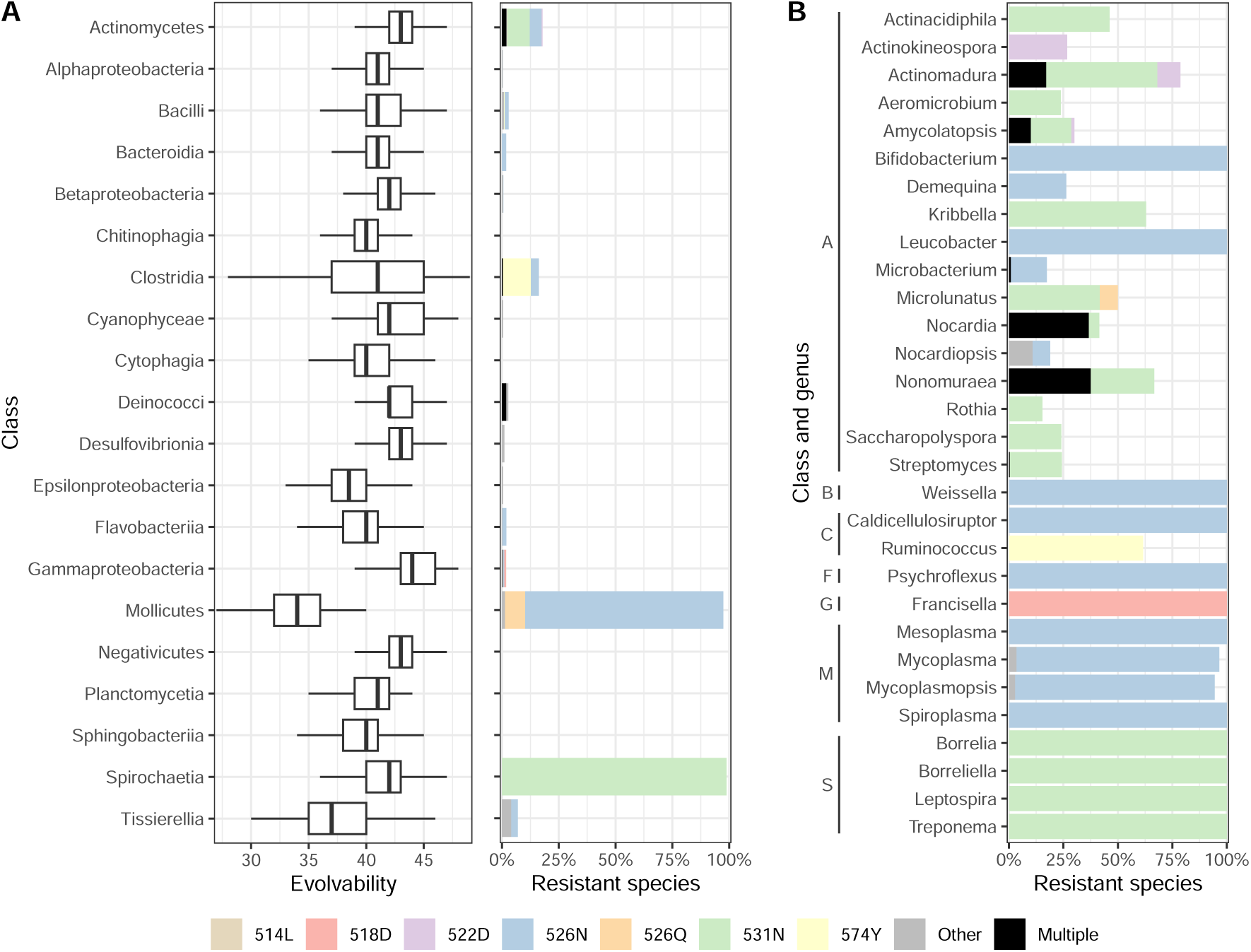
Predicted evolvability and intrinsic resistance at different taxonomic levels. The left plot in panel A shows the distribution of evolvability across the twenty most specious bacterial classes. Evolvability is defined as the total number of one-step single nucleotide changes that can result in amino acid substitutions (within our panel). The right-hand plot in panel A shows the percentage of species predicted to be intrinsically resistant in the same classes, with colours indicating which resistance mutation they carry. Only the six most common mutations are shown explicitly, all other mutations are shown in grey. Panel B shows the percentage of resistant species across the thirty genera with the highest percentage of resistant species. Only genera with at least twelve species were included. Genera are grouped by class, with the following abbreviations: A=Actinomycetes, B=Bacilli, C=Clostridia, F=Flavobacteriia, G=Gammaproteobacteria, M=Mollicutes, S=Spirochaetia.

There were 90 species with two copies of the *rpoB* gene in our dataset (most prominently belonging to genera *Nocardia*, *Nonomuraea* and *Actinomadura*; one species, *Oceanobacillus jeddahense*, had three copies). Sequence alignment revealed that in some species the two se-quences are identical at the main resistance-determining region of *rpoB* (Figure S4), indicating relatively recent duplication events (or potentially erroneous genome assemblies). However, most pairs of *rpoB* copies exhibit considerable divergence, and in most cases one copy carries one or several resistance mutations whereas the other one does not (Figure S4).

### The mutational spectrum of intrinsic and evolvable RIF resistance

Although overall evolvability of RIF resistance varies relatively little across bacterial groups, there is considerable variation when individual mutations are considered (Figure 4). Some mutations can readily arise in essentially all bacteria, either through one, two or even three different point mutations. Other mutations can arise in only some but not other species, and some mutations can only arise through a single point mutation in very few species. This variation also persists when mutations are pooled into amino acid positions, with some positions (e.g., 513 and 516) exhibiting consistently high evolvability whereas at other positions (e.g., 522 and 574) no mutation is evolvable in all or even the majority of species. Figure 4 also highlights which resistance mutations are already present in some species, the most common being 531N (565 species), 526N (546 species), and 574Y (183 species). Stratifying this mutational spectrum taxonomically demonstrates that heterogeneity in evolvability can differ considerably across classes (Figure S3). For example, mutation 509R can evolve via three different point mutations in most members of the Deinococci but this same mutation cannot evolve through a single point mutation in many other classes.

**Figure 4:**
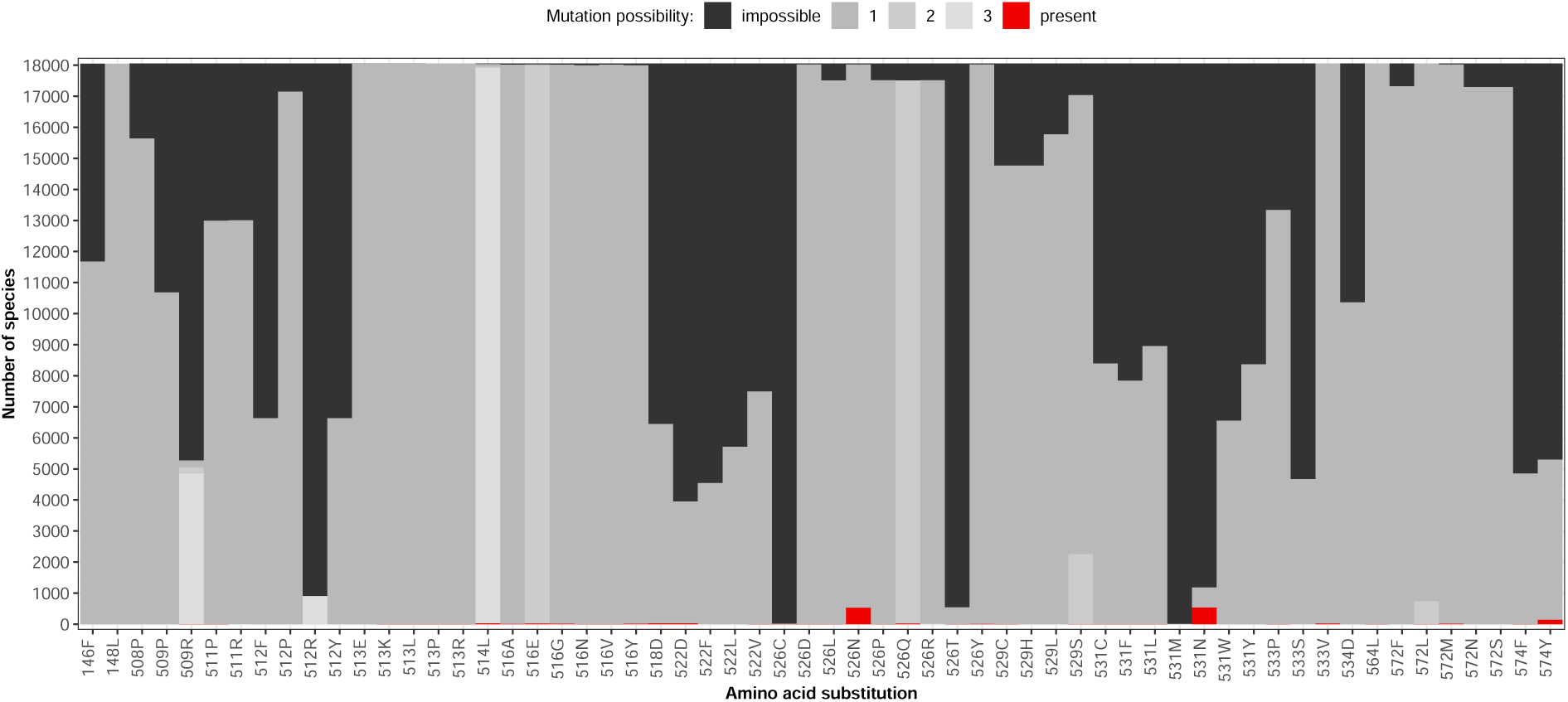
The mutational spectrum of intrinsic and evolvable RIF resistance. For each of the 60 amino acid residues in our panel of screened RIF resistance mutations, the plot shows the number of species in which this mutation is either present (red), evolvable through a single point mutation (grey), or impossible to evolve through a single point mutation (black). Evolvability is further distinguished by different shades of grey according to whether the amino acid mutation can arise through one, two or three different point mutations at the respective codon. Species with multiple *rpoB* sequences were excluded from this plot.

We can shed light on this variation in evolvability by considering mutational codon networks, i.e. networks at a specific site where only codons that differ by a single nucleotide (and thus are evolvable from each other) are connected. Figure 5 shows an example for position 534. At this position, a single amino acid residue (aspartic acid) has been reported to confer RIF resistance (534D). All species in our dataset carry a glycine (G) residue at this position, but all four codons coding for glycine are used across species (at frequencies of 26%, 40%, 17% and 17% for GGA, GGC, GGG and GGT, respectively). Only the GGC and the GGT codons can mutate towards codons coding for aspartic acid in a single step, which explains why many species (43%) cannot evolve resistance at position 534. A more complex scenario, at position 531, is shown in Figure S5. At this position, almost all species (96%) carry a serine residue. How many and which of the seven reported resistance mutations (531C, F, L, M, N, W and Y) can evolve in a given species depends on which of the six codons coding for serine is employed. For example, 531W can only evolve in species with a TCG codon, whereas 531F and 531Y can only evolve in species with a TCC or TCT codon.

**Figure 5:**
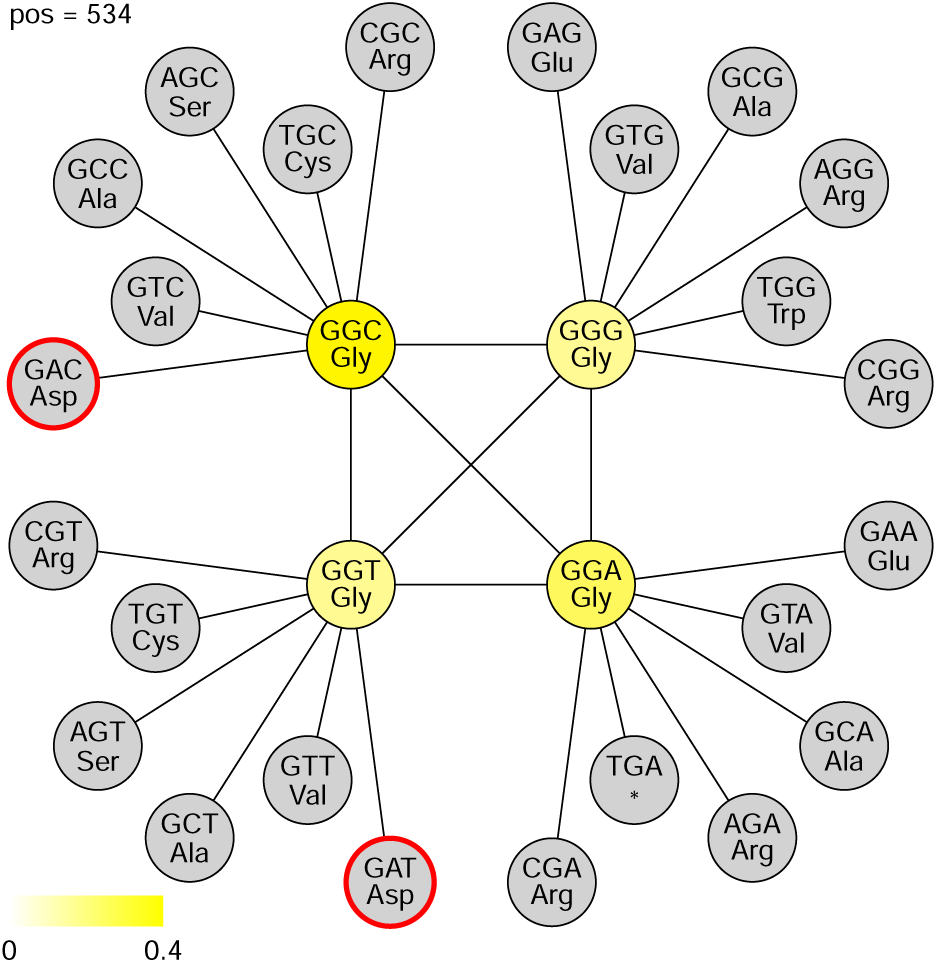
Incomplete codon network at *rpoB* amino acid position 534. Circles represent codons and lines connect codons that are separated by a single nucleotide difference. The yellow colour scale indicates the fraction of species that have a particular codon, with grey indicating that no species has the respective codon. Red circles indicate amino acids residues that have been reported to confer RIF resistance. For clarity, edges between unoccupied (grey) codons are not shown.

### Cross-validation with a high-throughput screen in *E. coli*

Finally, we sought to complement the results derived from our panel of reported mutations with data from a recent high-throughput screen by Yang *et al.*^11^. In this study, the authors employed multiplex automated genome engineering^20^ to create all possible single amino acid residues at the RIF binding site of *rpoB*, and through bulk competition estimated conferred RIF resistance for each mutation. As expected, there is substantial overlap between our panel of reported mutations and the mutations in Yang *et al.* (Figure S6). However, there are also many mutations not in our panel that Yang *et al.* predict to confer resistance. Conversely and more surprisingly, there are also many mutations that have been reported to confer resistance but do not show up as resistant in Yang *et al.* (Some but not all mutations in the latter group are at positions 146, 148, 508, 509 and 574 that were outside of the region screened in Yang *et al.*^11^.)

Figure S7 shows that the resistance mutations predicted in Yang *et al.*^11^ that are absent from our panel of reported mutations are generally characterised by low evolvability. With few exceptions, most of these amino acid residues cannot arise via a single point mutation, which explains why they have not been reported in previous mutant screens. Moreover, with one exception (525R), these mutations are either completely absent or very rare (*<* 0.2% frequency) within the species we screened. These results indicate that although there are likely many more resistance mutations theoretically possible than previously reported in mutant screens and natural isolates, these mutations are neither common in nature nor can they arise easily from existing genetic variation.

## Discussion

Predicting antibiotic resistance, a key challenge in medicine, public health and agriculture, requires knowledge of both selective pressures (driven mostly by antibiotic use) and evolvability (the amount of genetic variation for resistance, or the ease by which such variation can be generated). Focusing on the latter, we have analysed *rpoB* sequences from more than 18 000 species, enabling us to identify, explain and predict patterns of past and future RIF resistance evolution across the bacterial tree of life. Our approach underscores both the power and the limitations of leveraging genetic information across taxa.

Our results indicate that overall evolvability of RIF resistance is relatively homogeneous across the bacterial tree of life. Through a single nucleotide substitution, most species have immediate evolutionary access to between 35 and 47 resistance mutations, out of the 60 mutations that we screened. This means that the genetic code imposes only minor constraints on RIF resistance evolution (see also Ref. 21), and any large differences in mutation rates towards RIF resistance across species are unlikely to be explained by variation in evolvability as defined here. However, despite the homogeneity in total evolvability, there is considerable variation at the level of many individual mutations.

This variation can be important because individual RIF resistance mutations, even at the same position, can have very different phenotypic effects. Examples include different degrees of reduced competitive ability (“fitness costs”) in *Riemerella anatipestifer* ^22^, a wide range of minimum inhibitory concentrations and pleiotropic heat-stress resistance induced by different mutations at position 572 in *E. coli* ^23^, and divergence in patterns of carbon usage in *Bacillus subtilis* mutants carrying different *rpoB* mutations at positions 513 and 526^24^. Moreover, differ-ent RIF resistance mutations can also differ in their epistatic interactions with other resistance mutations^25,26^. For instance, whilst both positive and negative epistasis in competitive ability has been reported between RIF resistance mutation 526L in *rpoB* and various streptomycin resistance mutations in *rpsL*, no such epistasis was reported for 526N^25^. Finally, different resistance mutations at the same position may also determine future evolvability in different ways, for example by determining possible trajectories towards increased resistance ^26^ or reduced fitness costs (i.e. compensatory evolution^27^). Qi *et al.* reported very efficient compensatory evolution for the costly H526R mutation in *Pseudomonas aeruginosa*, but only incomplete compensatory evolution for H526Y^28^.

Around 8% of the species we screened carry at least one RIF resistance mutation. On this basis, we predict that several high-level taxa (e.g., Spirochaetia, Bifidobacterales, Mollicutes and Erysipelotrichales) as well as many smaller taxa are comprised entirely or predominantly of species that are naturally RIF resistant. RIF resistance mutations are likely to have occurred in the ancestors of many of these lineages in response to chemical compounds in their habitat that are structurally related to RIF. Similar to other drug-like metabolites, such chemicals are naturally released by bacteria to harm competitors, selecting for resistance in both producers and their targets^29–31^. RIF is a semisynthetic antibiotic that belongs to the rifamycin class of antibiotics. It is a derivative of rifamycin B, which was discovered in *Amycolatopsis rifamycinica* (then classified as *Streptomyces mediterranei*)^15^. Not surprisingly then, *A. rifamycinica* carries two RIF resistance mutations in *rpoB* (522D and 531N), and in fact around one third of the species in this genus carry one or both of these mutations. Related compounds belonging to the larger group of ansamycins that also exhibit antibacterial activities by inhibition of RNA polymerases are produced by other soil bacteria such as *Streptomyces* spp. and *Micromonospora halophytica*, as well as some marine and plant-associated bacteria^32^. Whilst it may be tempting to assume that all species carrying RIF resistance mutations evolved them in response to the presence of RIF-like compounds in their environment, these mutations could also have evolved in response to other selective pressures such as temperature^23^, or neutrally.

It is important to stress that the presence of a known, RIF resistance conferring mutation in a species does not inevitably result in phenotypic resistance. For example, 526N is ubiquitous in the Erysipelotrichales and Bifidobacterales, thus predicting natural RIF resistance in these groups. In accordance with this prediction, several studies have reported an increase in the frequency of members of the Erysipelotrichales in gut microbiota under RIF treatment^33,34^, but a corresponding increase in frequency has not been observed for species of *Bifidobacterium* ^33^. Similarly, whilst our prediction of RIF resistance in *Thermotoga maritima* (carrying mutation 533V) has independent experimental support^35^, *Francisella tularensis* appears to be susceptible to RIF despite harbouring resistance mutation 518D^36^. These examples indicate that at least some of the known RIF resistance mutations can become conditionally inactive through epistatic interactions, either with other sites within *rpoB* or with other genes. Investigating the extent and nature of these epistatic interactions, including careful experimental validation, will be an important task for the future.

One of the most notable *rpoB* mutations on which our investigation sheds light is 531N (as-paragine). This mutation has previously been reported to be responsible for intrinsic resistance in several *Streptomyces* ^37,38^ and some spirochaete species^39–41^. Our analysis shows that this mutation is the most frequent intrinsic resistance mutation, being present in 561 species (3.1% of all species in our dataset). In particular, 531N is universally present in the Spirochaetia and in a diverse range of species within the Actinomycetia. Surprisingly though, 531N is the only mutation in our panel that has not been reported in a mutant screen (only in natural isolates). We can explain this seemingly paradoxical result by our finding that evolvability of 531N is exceptionally low (Figure 4). The majority of species (*>* 93%) carry a TCA, TCC, TCT, or TCG codon at position 531, all coding for serine. These codons are more than one mutational step away from the two codons coding for asparagine (AAC or AAT). How then has 531N evolved in so many species? We speculate that there could be two major pathways enabling species to evolve the 531N residue. First, some species (*≈* 3%) have codon AGC or AGT (also coding for serine) at this position, from which an asparagine-coding codon can evolve in a single mutational step. Second, a species could first evolve a tyrosine residue (531Y, codon TAT or TAC) and from there, in a second mutational step, evolve 531N. 531Y is a resistance mutation reported in several mutant screens^10,24,37,42–46^ but in our screen is not predicted to be intrinsically present in any species. It appears then that 531N is a mutation that is hard to evolve for most species but can confer strong resistance without imposing strong pleiotropic fitness costs, relative to easier-to-evolve mutations.

Several caveats and limitations in our approach merit consideration. First, our panel of RIF resistance mutations involves an inherent trade-off between comprehensiveness (including as many reported mutations as possible) and robustness (ensuring that no erroneous or very species-specific mutations are included). Our cross-validation with an independent data set^11^ indicates that our filtering options navigated this trade-off reasonably well. Second, we only considered point mutations in *rpoB*. Both insertion and deletion mutations in *rpoB* have been reported to confer resistance to RIF as well^7,10^. However, these mutations tend to be rarer and less repeatable than point mutations, so that assessing their evolvability would be very hard. Resistance to RIF can also be caused by other mechanisms, including various enzymes that deactivate RIF, efflux pumps (reviewed in Ref. 16) and HelR proteins that dissociate RIF from the RNA polymerase (reviewed in Ref. 47). Third, we have employed deliberately simple, binary measures of evolvability in our study, quantifying the number of mutations that can arise through a single nucleotide change and the number of nucleotide changes producing a mutation. A more fine-grained measure would also include differences in mutation rates at the nucleotide level, in particular accounting for higher transition than transversion mutations as well as higher mutation rates at specific sites^48,49^. However, including such differences would still not capture differences in baseline mutation rates across species, which are largely unknown. Finally, we have (with very few exceptions) only considered a single genome per species. This means that there may be species not predicted to be resistant to RIF that are in fact polymorphic for RIF resistance. This is trivially true for many pathogens that are exposed to RIF but may also be true for non-pathogenic bacteria. In fact, our finding that intrinsic RIF resistance in the genus *Streptomyces* exhibits little phylogenetic clustering (Figure S2) could be explained by widespread but polymorphic resistance in this group, where type strains with deposited genomes sometimes do and sometimes do not exhibit RIF resistance.

In conclusion, our study provides a phylogenetically broad characterisation of the mutational landscape of resistance to RIF, an antibiotic of major clinical significance. The approach and bioinformatics pipeline we developed can readily be extended to other antibiotics where resis-tance is acquired through point mutations in highly conserved genes, including aminoglycosides and quinolones. Pending experimental validation, our predictions on intrinsic resistance and evolvability of RIF resistance could help assess the impact of RIF on commensal gut bacteria as well as assist in the design of probiotics as an alternative therapeutic strategy to eliminate antibiotic resistant pathogens.

## Acknowledgments

We thank Mary Gannon for help surveying the literature on rifampicin resistance, and acknowl-edge funding from the Australian research council to JE (grant number DP190102485).

## Methods

### Selection of mutation panel

We sought to assemble a panel of mutations within the gene encoding the beta subunit of RNA polymerase, *rpoB*, that have been reported to confer resistance to RIF. To this end, we compiled studies which reported single amino acid substitutions resulting in phenotypic resistance to RIF, obtained either experimentally (mutant screens or genetic constructs), or in clinical or environmental isolates. We used Google Scholar to find relevant studies using the following key words: “rifampicin resistance”, “RNA polymerase subunit beta”, “point mutation” and “amino acid substitution”. The following information was then extracted from the obtained studies: species name, origin (experimental studies vs. natural or clinical isolates), as well as mutated amino acids and their corresponding positions. Standardised amino acid positions in *E. coli* coordinates were obtained by alignment of the respective *rpoB* sequence to the *rpoB* reference sequence in (*E. coli* MG1655). To avoid the inadvertent inclusion of incidental mutations which may not robustly reflect the evolution of RIF resistance, we only considered mutations reported in at least three studies and within at least three different species.

### Prediction of presence and evolvability of rifampicin resistance conferring mutations

We next determined the existence and evolvability of mutations in our panel across all bacterial species with published genomes. First, we downloaded reference and/or representative genomes at all assembly levels (contig, scaffold and complete) from the NCBI Genome database^50^. The *rpoB* sequence of each genome was extracted from their annotated genomic coding sequences using the following keywords: “rpoB”, “DNA-directed RNA polymerase subunit beta”, “DNA-directed RNA polymerase subunit beta chain”, “RNA polymerase, beta subunit”, “DNA-directed RNA polymerase subunit beta”. Only *rpoB* sequences with a length of at least 3000 base pairs were retained.

To further ensure the robustness of these *rpoB* sequences, we translated and aligned them to the *E. coli* MG1655 reference *rpoB* sequence, extracted the amino acid sequence from positions 501 to 600 (the ‘core’ region encompassing the vast majority of RIF resistance mutations). We calculated the Levenshtein distance^51^ between the core sequence region of the focal sequence to the corresponding homologous region of the *E. coli* MG1655 reference sequence. Sequences with a distance greater than 35 were discarded. In total, 444 out of 18682 *rpoB* sequences were discarded due to the length or Levenshtein distance filter.

The filtered sequences were then screened for the presence and evolvability of mutations in our panel of reported mutations. Specifically, for each combination of a reported resistance-conferring amino acid residue and an *rpoB* sequence, we determined: 1) whether the mutation is already present; and 2) in how many different ways (including zero) the mutation could evolve through a one-step single nucleotide change given the codon at the respective position with the *rpoB* sequence. We summarised this data at the level of *rpoB* sequences to determine overall predicted resistance (at least one resistance mutation present) and overall evolvability (total number of mutations that can evolve in a single mutational step).

### Phylogenetic analyses

In order to investigate the phylogenetic distribution of naturally RIF resistant species as well the their evolvability, we downloaded a large phylogenetic tree of all bacterial groups from the Genome Taxonomy Database (GTDB; https://gtdb.ecogenomic.org/)^19^. The GTDB tree is based on genome trees constructed using an aligned set of 120 single bacterial protein markers as well as ribosomal proteins and ribosomal RNA genes. The genomes were reference sequences submitted in NCBI (starting with release 1976)^50^ and were independently quality controlled using CheckM^52^. We subset the GTDB tree for species included in our final screen. We investigated phylogenetic clustering of resistance and evolvability by calculating Blomberg’s *K* and Pagel’s *λ* statistics^53,54^.

### Cross-validation using an alternative panel of experimentally predicted rifampicin resistance in *E. coli*

We corroborated our predictions with a secondary mutational screen using a mutation panel employed by Yang *et al.*^11^. The authors of this article constructed a comprehensive set of *E. coli* mutants (760 mutants) carrying all possible single amino acid substitutions within *rpoB* at positions 510—537 and 563—572 to investigate the effects of *rpoB* mutations on the efficacy of RIF binding. From these 760 mutations, we selected those that increased in frequency in RIF-containing medium (minimum log2(fold-change) of zero), exempting those mutations that

were already included in our own mutation panel. We then screened all *rpoB* sequences for the presence and evolvability of these mutations, in the same way that we did for the mutations in our panel of reported mutations.

### Implementation

All analyses were conducted in R version 4.4.1^55^. Bioinformatics analyses were performed us-ing packages rentrez v1.2.3^56^, taxonomizr v0.10.6^57^, Biostrings v2.58.0^58^, pwalign v1.0.0^59^, stringdist v0.9.12^60^, future.apply v1.11.2^61^. Packages ape v5.8.0^62^, castor v1.8.0^63^, ggtree v4.4.0^64^, tidytree v0.4.6^65^ and treeio v1.29.0^66^ were used for phylogenetic analysis and tree vi-sualisation. Data wrangling and visualisation were performed using packages tidyverse v2.0.0^67^, ggnewscale v0.4.10^68^, GGally v2.2.1^69^, ggpubr v0.6.0^70^, ggh4x v0.2.8^71^, RColorBrewer v1.1-3^72^ and patchwork v1.2.0^73^. All code for this project will be made available as a GitHub release upon publication.

## Supplementary figures

**Figure S1:**
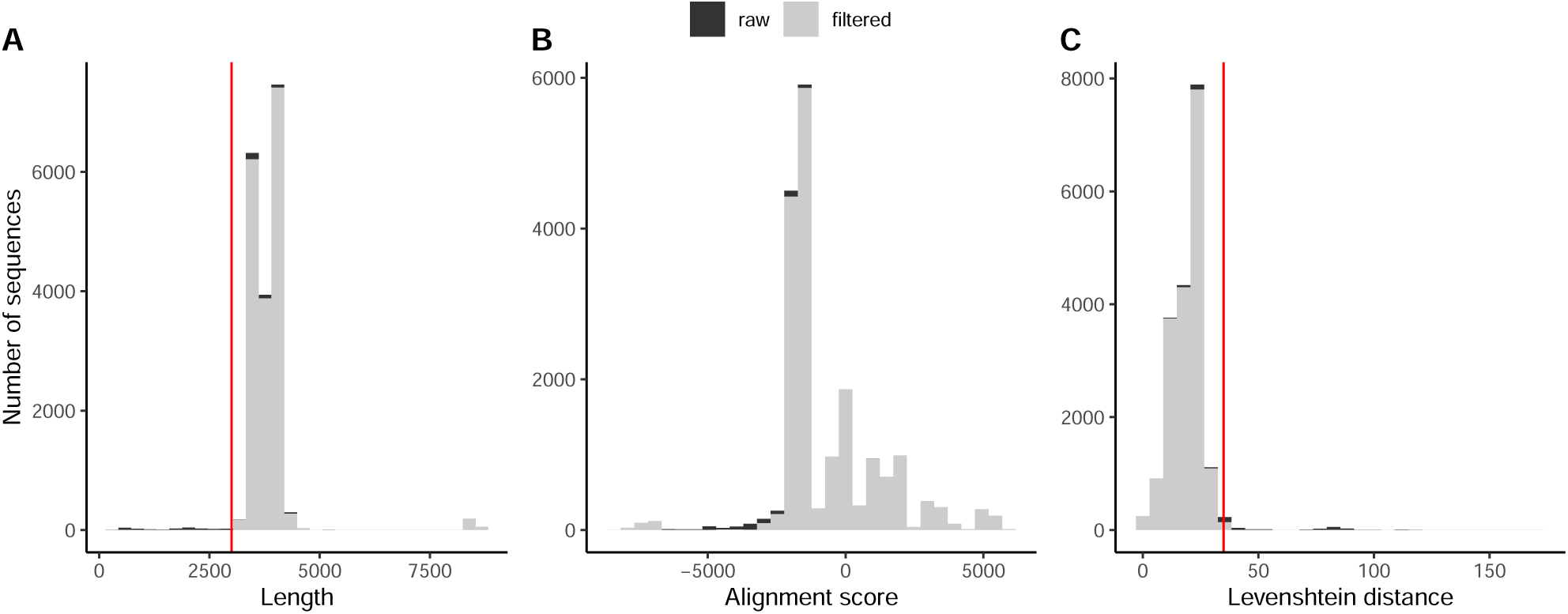
Properties of *rpoB* sequences extracted from genomes downloaded from NCBI. A total of 18 682 sequences were extracted from 18 469 genomes. The plots show the distribution of properties of the extracted *rpoB* sequences, with black colour indicating the distribution before and grey the distribution after filtering. Panel A) shows the distribution in the length of the *rpoB* sequences, B) the alignment score of each sequence to the *E. coli rpoB* reference sequence, and C) the Levenshtein distance of the core region of the sequence (between amino acid position 500 and 600) to the *E. coli rpoB* reference sequence. In A) and C), the vertical red lines indicate the applied cut-offs used for filtering (minimum length of 3000 and maximum Levenshtein distance of 35, respectively).

**Figure S2:**
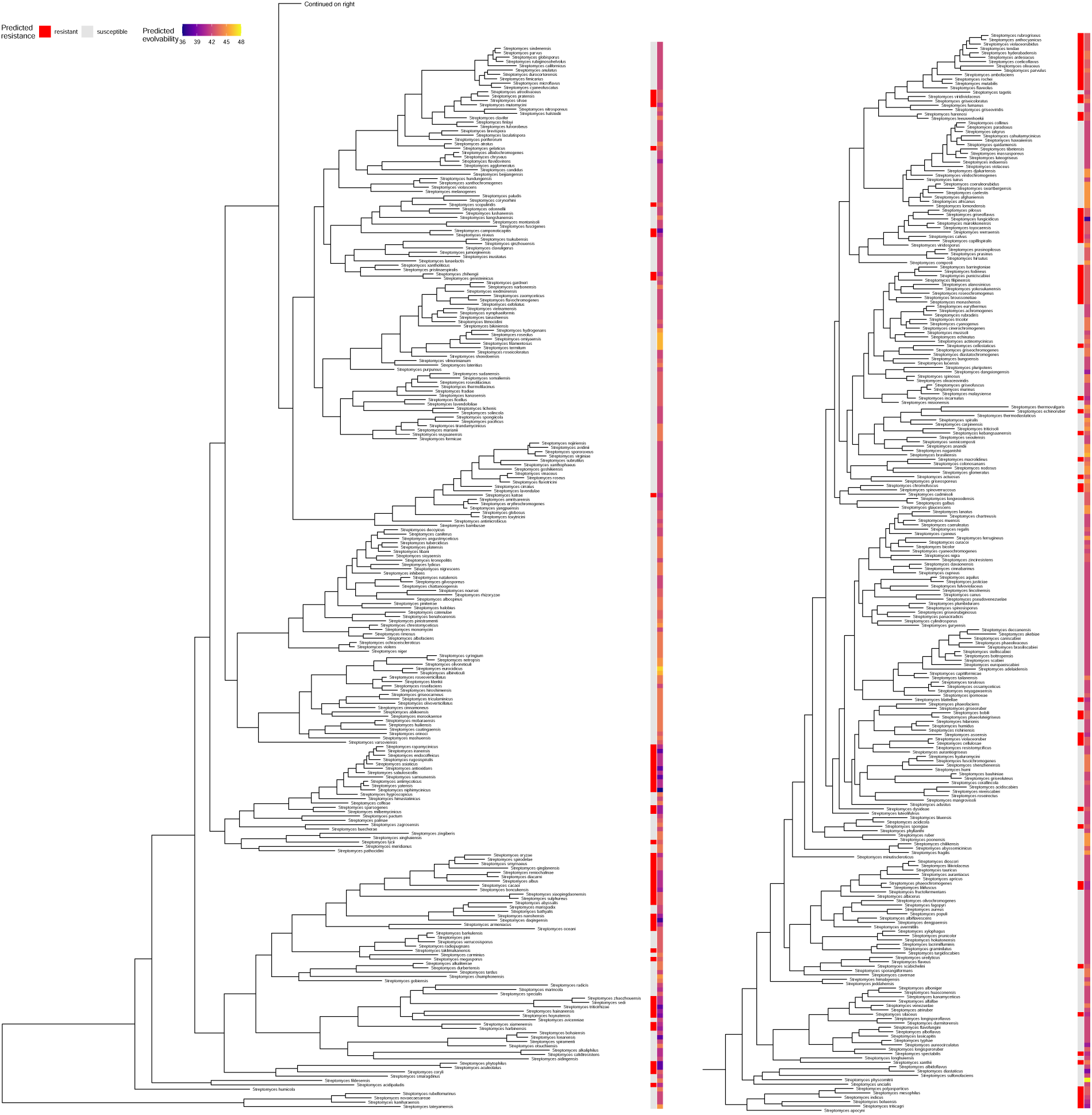
Phylogenetic tree of the genus *Streptomyces*, obtained as a subtree of the tree shown in Figure 2. Predicted intrinsic resistance and evolvability are indicated in the same way as in Figure 2. The tree is shown in two parts, with the top-most branch of the subtree on the left-hand side connecting to the subtree on the right.

**Figure S3:**
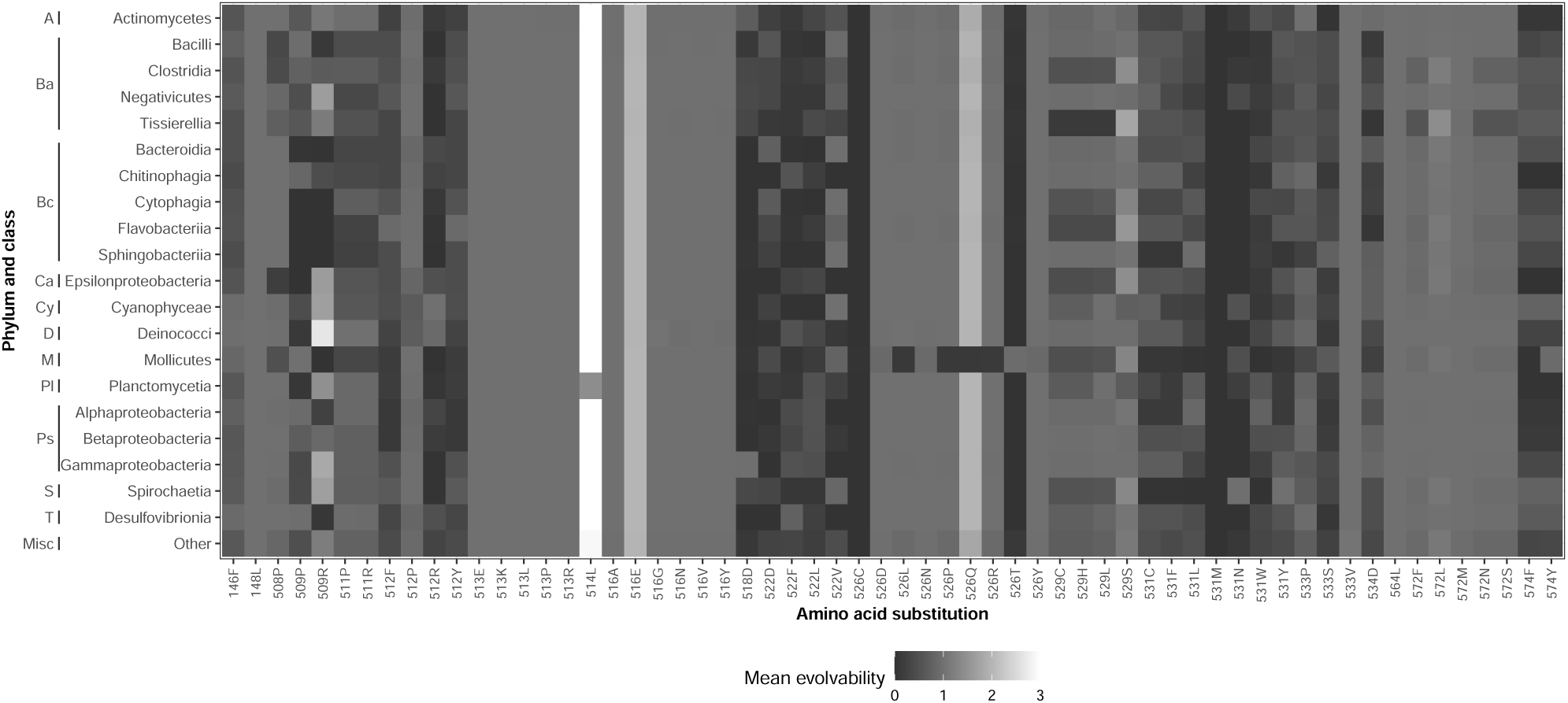
Mean evolvability by bacterial class and mutation. Here, evolvability is calculated as the mean number of nucleotide substitutions across species that would produce any given amino acid residue. Bacterial classes are grouped by phylum, with the following abbreviations used: A=Actinomycetota, Ba=Bacillota, Bc=Bacteroidota, Ca=Campylobacterota, Cy=Cyanobacteriota, D=Deinococcota, M=Mycoplasmatota, Pl=Planctomycetota, Ps=Pseudomonadota, S=Spirochaetota, and T=Thermodesulfobacteriota. Only the 20 most specious classes are shown.

**Figure S4:**
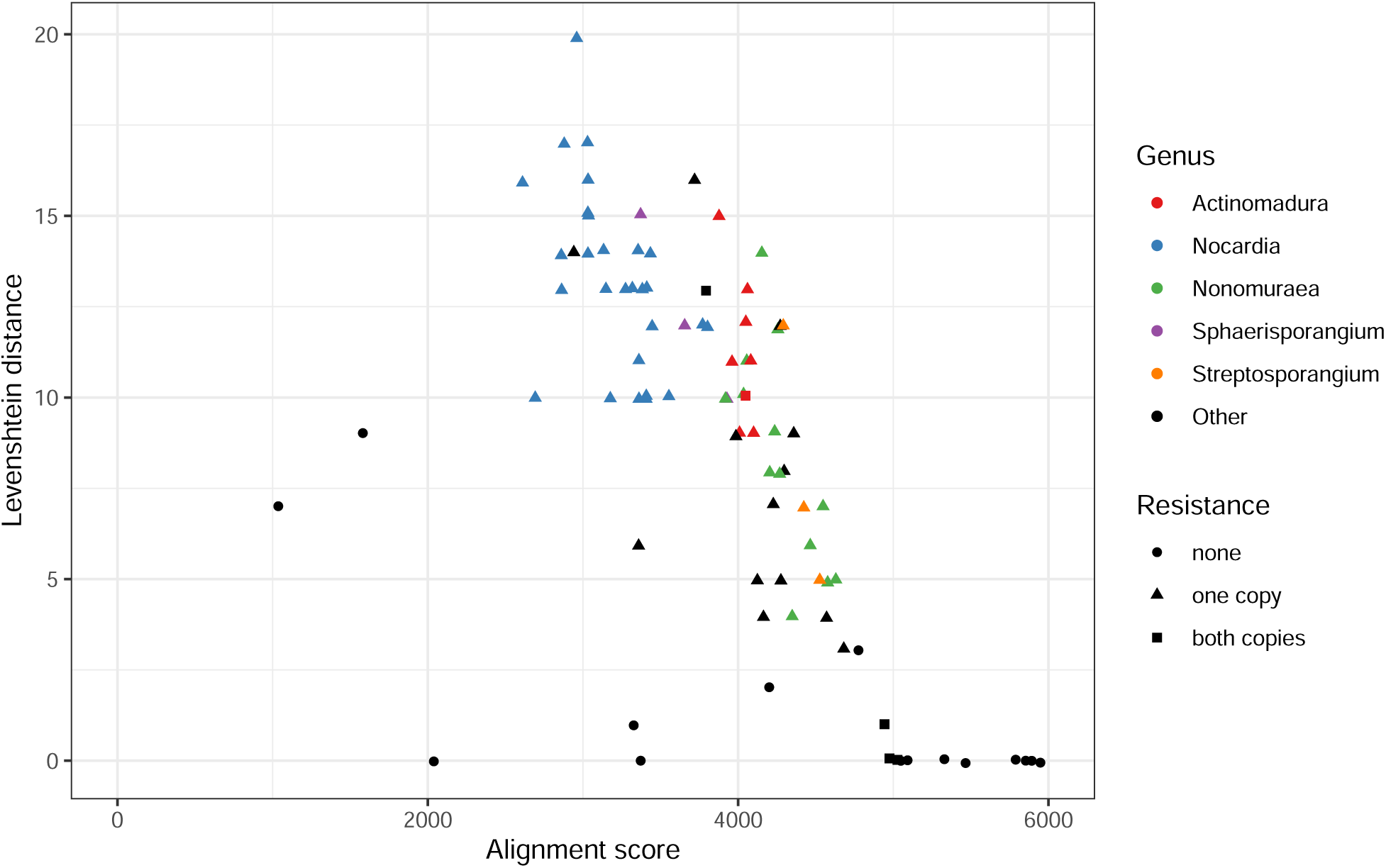
Comparison of pairs of *rpoB* sequences across the 90 species in our dataset that had two *rpoB* copies. For each pair, the *rpoB* sequences were aligned and the plot shows both alignment scores and Levenshtein distance between the core *rpoB* regions (positions 501 through 600). Pairs of sequences are coloured by genus and shapes show their resistance status, i.e., whether one, both or none of the copies are predicted to confer RIF resistance.

**Figure S5:**
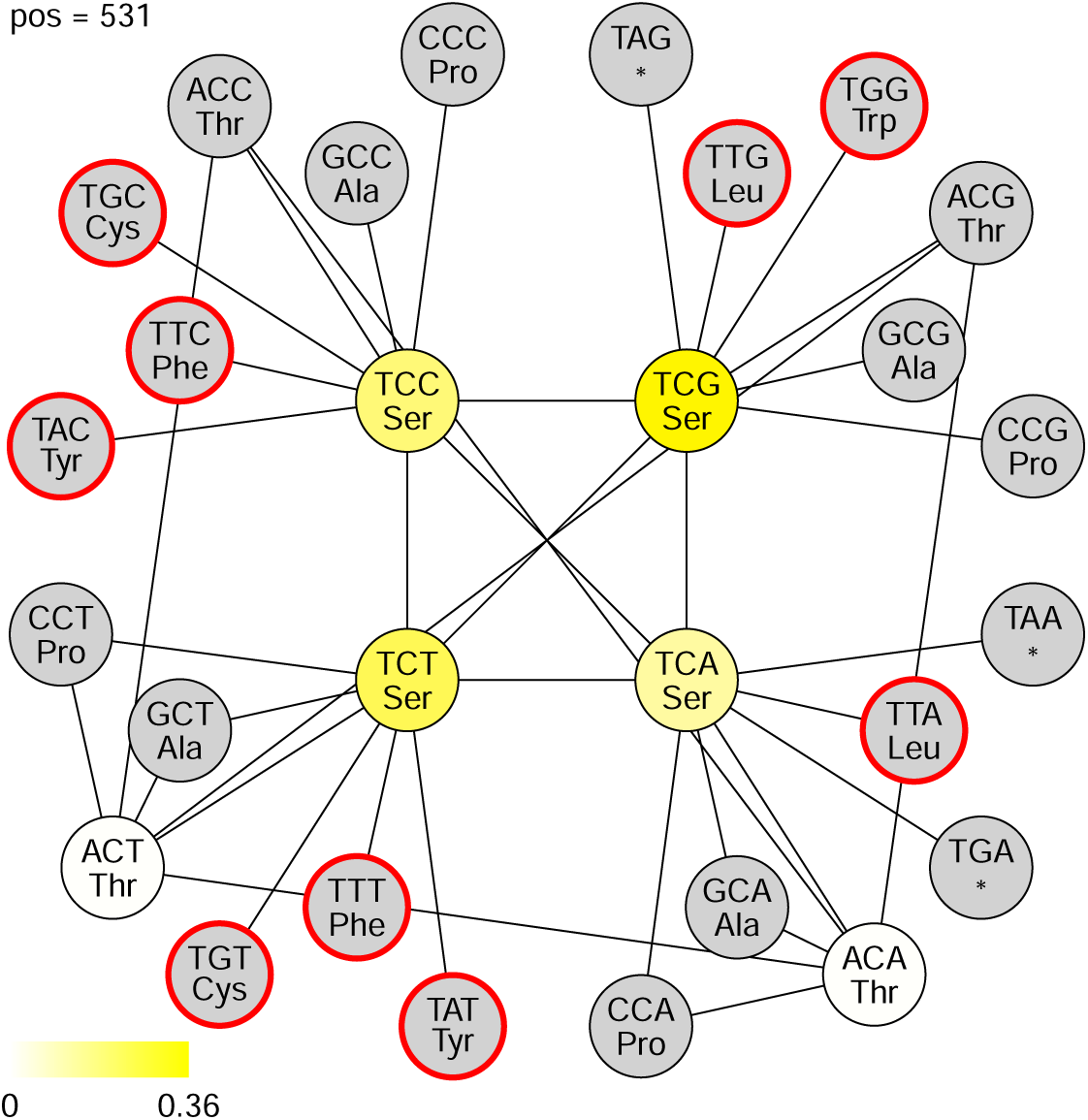
Incomplete codon network at *rpoB* amino acid position 531. Circles represent codons and lines connect codons that are separated by a single nucleotide difference. The yellow colour scale indicates the fraction of species that have a particular codon, with grey indicating that no species has the respective codon. Red circles indicate amino acids residues that have been reported to confer RIF resistance. For clarity, edges between unoccupied (grey) codons are not shown. Note that although most species (*≈* 93%) have one of the six codons shown in yellow or white in this figure, many species have other codons that for simplicity are not shown in this figure.

**Figure S6:**
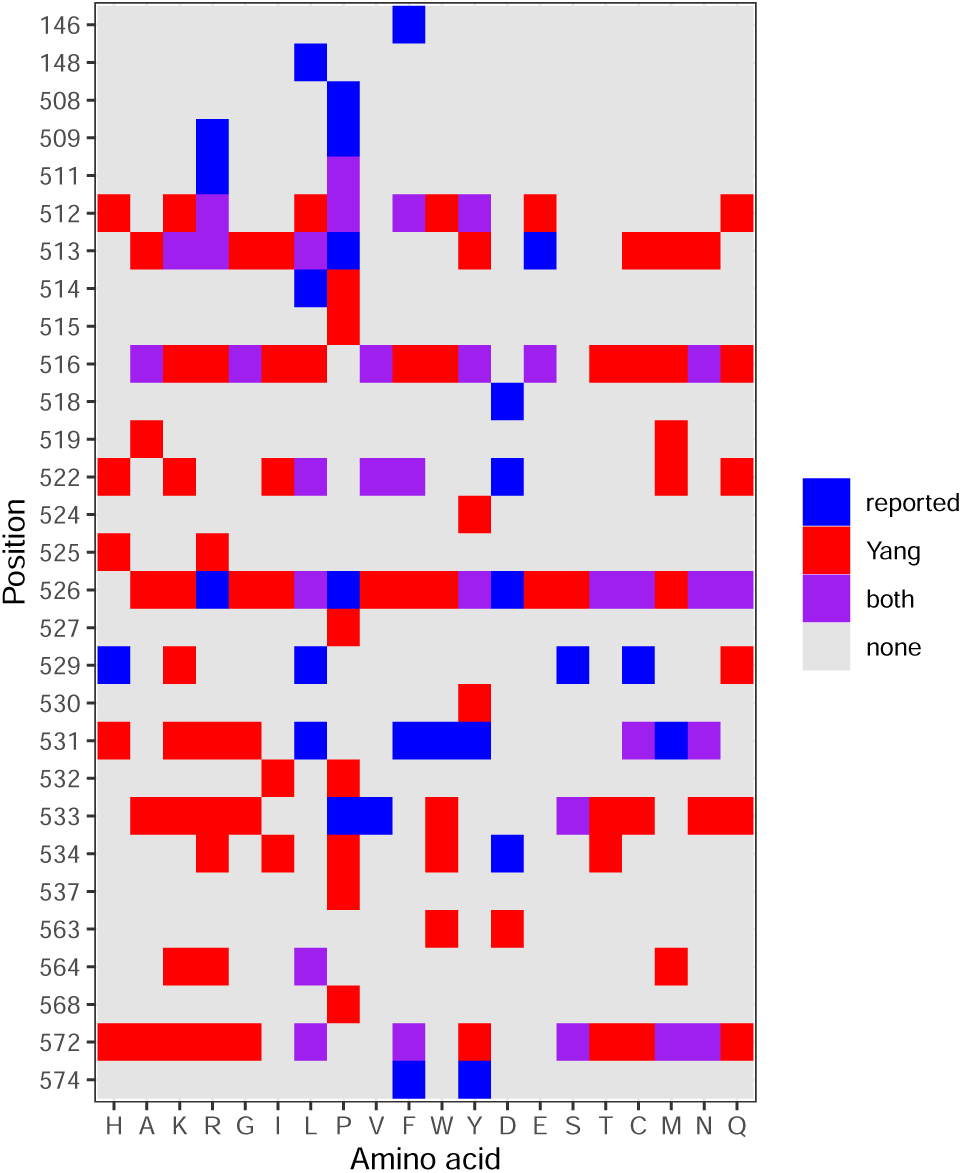
Comparison between the reported mutations in our screened panel, and the mutations predicted to confer RIF resistance in the high-resolution screen reported by Yang *et al.* ^11^. Each square represents an amino acid residue across a range of positions within the *rpoB* gene. Blue indicates mutations in our panel that are not reported in Yang *et al.* ^11^, red indicates mutations reported in Yang *et al.* ^11^ but absent from our panel, purple indicates mutations in both, and grey indicates amino acid residues not predicted to confer RIF resistance in either set of mutations.

**Figure S7:**
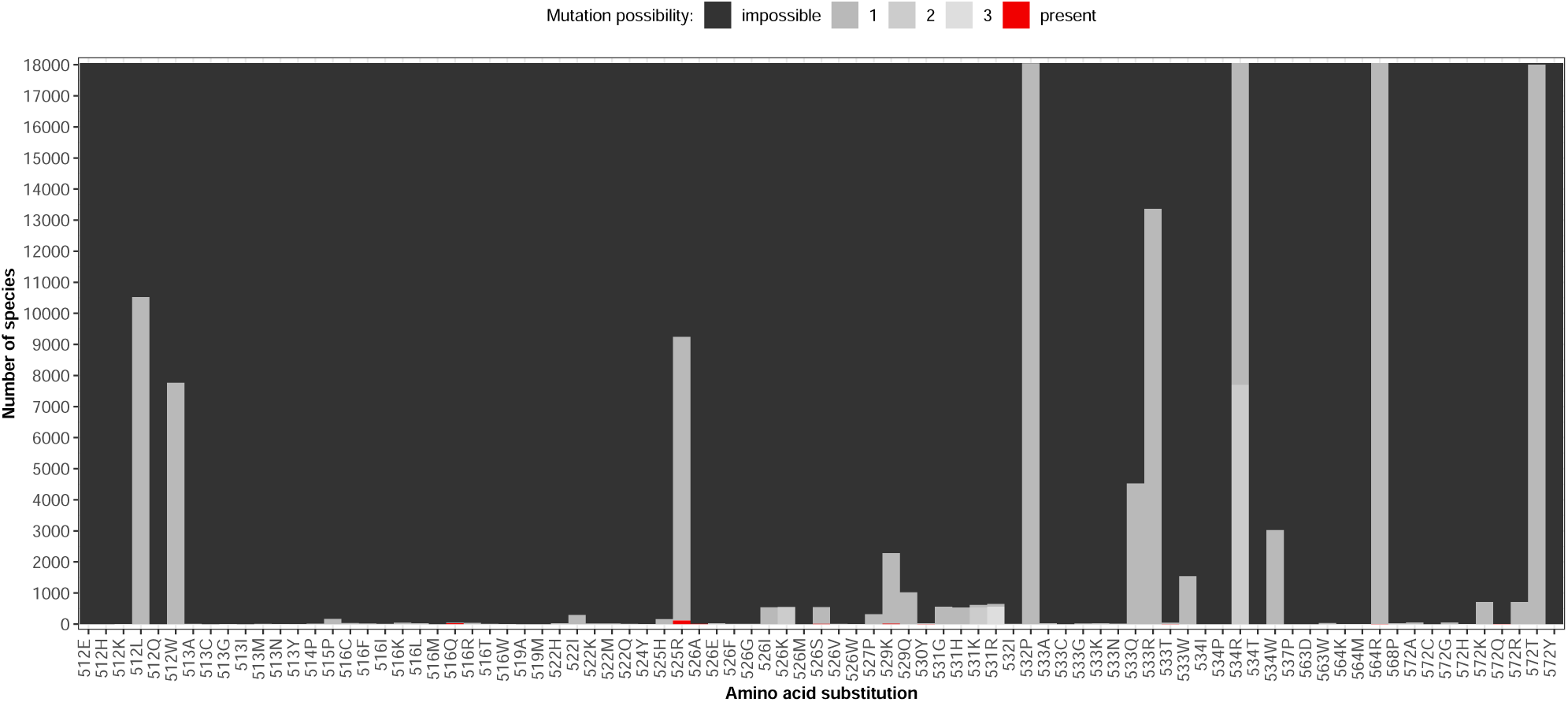
The mutational spectrum of intrinsic and evolvable RIF resistance for mutations predicted to confer RIF resistance in Yang *et al.* ^11^ that were not present in our panel of reported RIF resistance mutations. As in Figure 4, the plot shows the number of species in which this mutation each either present (red), evolvable through a single point mutation (grey), or impossible to evolve through a single point mutation (black). Evolvability is further distinguished by different shades of grey according to whether the amino acid mutation can arise through one, two or three different point mutations at the respective codon. Species with multiple *rpoB* sequences were excluded from this plot.

